## Supplementary material for "Modelling interference between vectors of non-persistently transmitted plant viruses to identify effective control strategies": S1 Text

### S1 Text. Generalized interference functions

The visiting and emigration interference functions presented in the main text for the direct and indirect interference scenarios (Eqs 4 and 5 in the main text) can be extended to include also intermediate interference scenarios via the following functions:

$$f(N_R) = \frac{1}{1 + \left( \frac{\nu_1}{h_R^{1-\beta_1}} \frac{N_R}{h^{\beta_1}} \right)^{\alpha_1}} \qquad g(N_R) = 1 + \left( \frac{\nu_2}{h_R^{1-\beta_2}} \frac{N_R}{h^{\beta_2}} \right)^{\alpha_2} \tag{1}$$

The parameter $\beta_1 \in [0,1]$ ($\beta_2 \in [0,1]$) allows different interference scenarios to be represented. When $\beta_1 = 1$ ($\beta_2 = 1$), the function $f(N_R)$ $[g(N_R)]$ represents direct interference, when $\beta_1 = 0$, the function $f(N_R)$ $[g(N_R)]$ represents indirect interference, when $0 < \beta_1 < 1$ the function $f(N_R)$ $[g(N_R)]$ represents forms of interference between these two extremes. The other parameters have the same interpretations as in the main text. Note that, in our model, the value of the "strength" parameter $\nu_1$ ($\nu_2$) is defined for the direct interference scenario ($\beta_1 = \beta_2 = 1$). To assure that its value is biologically relevant also for other interference scenarios, we scale it with $h_R^{1-\beta_1}$ ($h_R^{1-\beta_2}$). This implies that, when $h = h_R$ - all other things being equal [*i.e.* the values of $\alpha_1$ ($\alpha_2$) and $N_R$] - the value of the interference function is the same independently from the underlying interference scenario.
