## Supplementary material for "Modelling interference between vectors of non-persistently transmitted plant viruses to identify effective control strategies": S2 Text

### S2 Text. Mathematical analysis of the single host-multi vector model

#### Non dimensionalization

The system of differential equations corresponding to the model, after integrating the assumption in the main text ($N_P$ constant and $\lambda = T\tau$), and after including the dynamics of the population size of resident ($N_R = X_R + Z_R$) and transient ($N_T = X_T + Z_T$) aphids, can be rewritten as

$$\begin{cases} \dot{I} = (\Lambda_R \delta_R Z_R + \Lambda_T \delta_T f(N_R) Z_T)(N_P - I) - (\rho + \theta) I \\ \dot{N_R} = r N_R (1 - \frac{N_R}{h}) - \mu N_R \\ \dot{Z_R} = \Lambda_R \varepsilon_R \frac{I}{N_P}(N_R - Z_R) - (\gamma + \mu) Z_R \\ \dot{N_T} = \tau (T - g(N_R) N_T) \\ \dot{Z_T} = \pi \tau T + \Lambda_T \varepsilon_T f(N_R) \frac{I}{N_P}(N_T - Z_T) - (\gamma + \tau g(N_R)) Z_T \end{cases} \tag{1}$$

Where

$$f(N_R) = \frac{1}{1 + \left( \frac{\nu_1}{h_R^{1-\beta_1}} \frac{N_R}{h^{\beta_1}} \right)^{\alpha_1}} \tag{2}$$

$$g(N_R) = 1 + \left( \frac{\nu_2}{h_R^{1-\beta_2}} \frac{N_R}{h^{\beta_2}} \right)^{\alpha_2} \tag{3}$$

Note that, as presented in the previous section, by considering $\beta_1 = 1$ ($\beta_1 = 0$) the visiting interference function $f(N_R)$ represents the direct (indirect) interference scenario presented in the main text. Similar considerations are taken for the emigration interference $g(N_R)$.

The equations of $\dot{S}$, $\dot{X}_R$ and $\dot{X}_T$ have been omitted because they can be derived from $S = N_P - I$, $X_R = N_R - Z_R$ and $X_T = N_T - Z_T$, respectively. To highlight the resident aphid carrying capacity, the equation for $N_R$ can be rewritten as

$$\dot{N_R} = r' \left( 1 - \frac{N_R}{\kappa} \right) \tag{4}$$

where

$$r' = r - \mu \qquad \kappa = h \left( 1 - \frac{\mu}{r} \right) \tag{5}$$

By making the transformation

$$
\begin{aligned}
I &= N_P \hat{I} && (6)\\
N_R &= \kappa \hat{N}_R && (7)\\
Z_R &= \kappa \hat{Z}_R && (8)\\
N_T &= T \hat{N}_T && (9)\\
Z_T &= T \hat{Z}_T && (10)\\
t &= \frac{\hat{t}}{\rho+\theta} && (11)
\end{aligned}
$$

and writing

$$
f(\hat{N}_R) = \frac{1}{1+\left(\frac{\nu_1}{\hat{h_R}^{1-\beta_1}}\frac{\hat{N}_R}{\hat{h}^{\beta_1}}\right)^{\alpha_1}} = f(\cdot) \tag{12}
$$

$$
g(\hat{N}_R) = 1+\left(\frac{\nu_2}{\hat{h_R}^{1-\beta_2}}\frac{\hat{N}_R}{\hat{h}^{\beta_2}}\right)^{\alpha_2} = g(\cdot) \tag{13}
$$

where $\hat{h}_R$, $\hat{h}_1$ and $\hat{h}_2$ are the non dimensional versions of the parameter $h_R$, $h_1$ and $h_2$, that is

$$
\begin{aligned}
h_R &= \kappa \hat{h}_R && (14)\\
h_1 &= \kappa \hat{h}_1 && (15)\\
h_2 &= \kappa \hat{h}_2 && (16)
\end{aligned}
$$

we obtained a nondimensionalized version of the original model equations:

$$
\begin{cases}
\dot{\hat{I}} = \left(i_R \hat{Z}_R + i_T f(\cdot)\hat{Z}_T\right)(1-\hat{I}) - \hat{I}\\
\dot{\hat{N}}_R = q\hat{N}_R\left(1-\hat{N}_R\right)\\
\dot{\hat{Z}}_R = a_R(\hat{N}_R - \hat{Z}_R)\hat{I} - m\hat{Z}_R\\
\dot{\hat{N}}_T = e(1-g(\cdot)\hat{N}_T)\\
\dot{\hat{Z}}_T = u + a_T f(\cdot)(\hat{N}_T - \hat{Z}_T)\hat{I} - (d+eg(\cdot))\hat{Z}_T
\end{cases} \tag{17}
$$

The composite parameters are given by

$$
i_R = \frac{\Lambda_R \delta_R \kappa}{\rho+\theta} \qquad i_T = \frac{\Lambda_T \delta_T T}{\rho+\theta} \qquad q = \frac{r'}{\rho+\theta} \tag{18}
$$

$$
a_R = \frac{\Lambda_R \varepsilon_R}{\rho+\theta} \qquad m = \frac{\gamma+\mu}{\rho+\theta} \qquad u = \frac{\pi\tau}{\rho+\theta} \tag{19}
$$

$$
a_T = \frac{\Lambda_T \varepsilon_T}{\rho+\theta} \qquad d = \frac{\gamma}{\rho+\theta} \qquad e = \frac{\tau}{\rho+\theta} \tag{20}
$$

#### Basic reproduction number

Considering the infected subsystem (*i.e.* the equations that describe the production of new infections and changes in state among infected individuals) of the nondimenional system in Eq 17, linearized around the infection-free steady state, and assuming that $u = 0$, which is equivalent to assuming that only non viruliferous transient aphids immigrate into the system, we determined the basic reproduction number of the disease, $R_0$, using the next generation method [2].

The matrix F describes the production of new infections and matrix the V describes changes in state.

$$F = \begin{bmatrix} 0 & i_R & i_T f(\cdot) \\ a_R \hat{N}_R & 0 & 0 \\ a_T f(\cdot) \hat{N}_T & 0 & 0 \end{bmatrix} \qquad V = \begin{bmatrix} -1 & 0 & 0 \\ 0 & -m & 0 \\ 0 & 0 & -(d + eg(\cdot)) \end{bmatrix} \tag{21}$$

$$F(-V)^{-1} = \begin{bmatrix} 0 & \frac{i_R}{m} & \frac{i_T f(\cdot)}{d+eg(\cdot)} \\ a_R \hat{N}_R & 0 & 0 \\ a_T f(\cdot) \hat{N}_T & 0 & 0 \end{bmatrix} \tag{22}$$

The basic reproduction number is the dominant eigenvalue ($\Lambda$) of the matrix $F(-V)^{-1}$, which can be easily computed by the characteristic equation of the matrix $F(-V)^{-1}$:

$$\Lambda^2 - \frac{i_R}{m} a_R \hat{N}_R - \frac{i_T f(\cdot)^2}{d + eg(\cdot)} a_T \hat{N}_T = 0 \tag{23}$$

Note that, from Eq 17, *i*) the value of $\hat{N}_R$ at the infection-free steady state is $\hat{N}_R = 1$ when $q > 0$ and $\hat{N}_R = 0$ when $q < 0$, and *ii*) the value of $\hat{N}_T$ at the infection-free steady state is $\hat{N}_T = \frac{1}{g(\cdot)}$.

$$R_0 = \frac{i_R a_R}{m} + \frac{i_T f(\cdot)^2 a_T}{g(\cdot)(d + eg(\cdot))} = R_0^R + R_0^T \qquad \text{if q} > 0 \tag{24}$$

$$R_0 = \frac{i_T f(\cdot)^2 a_T}{g(\cdot)(d + eg(\cdot))} = R_0^T \qquad \text{if q} < 0 \tag{25}$$

where $R_0^R$ corresponds to the virus transmission by resident aphids and $R_0^T$ considers the virus transmission by transient aphids. It is important to note that the quantities identified in Eqs 24 and 25 are actually $R_0^2$, since two cycles are involved in transmission, *i.e.* from plant to vector and from vector to plant [1]. However, since the two thresholds predict identical behaviour in terms of disease invasion (the threshold $R_0 = 1$ is precisely equivalent to $R_0^2 = 1$), we prefer to use the simpler formulation in our work.

The basic reproduction number can be rewritten with biological parameters, considering that at the infection-free steady state the value of $N_R = \kappa$ if $r > \mu$, or $N_R = 0$, otherwise, and the value of $N_T = \frac{T}{g(\kappa)}$. The interference functions $f(\cdot)$ and $g(\cdot)$ are equal to 1 if $N_R = 0$.

$$R_0 = \frac{1}{\rho+\theta}\left(\frac{\Lambda_R^2 \delta_R \varepsilon_R \kappa}{\gamma+\mu} + \frac{\Lambda_T^2 \delta_T \varepsilon_T f(\kappa)^2 T}{g(\kappa)(\gamma+\tau g(\kappa))}\right) \text{ if } r > \mu \tag{26}$$

$$R_0 = \frac{\Lambda_T^2 \delta_T \varepsilon_T T}{(\rho+\theta)(\gamma+\tau)} \text{ if } r < \mu \tag{27}$$

### Epidemic equilibrium

The equilibrium values of $\hat{N}_R$ come from a solution of

$$0 = q\hat{N}_R(1-\hat{N}_R) \tag{28}$$

When $q = r - \mu < 0$, the equilibrium that is stable in the long term is $\hat{N}_R = 0$, when $q = r - \mu \geq 0$ the equilibrium that is stable in the long term is $\hat{N}_R = 1$. The equilibrium value of $\hat{N}_T$ comes from the solution of

$$0 = e(1 - g(\cdot)\hat{N}_T) \tag{29}$$

$$\hat{N}_T = \frac{1}{g(\cdot)} \tag{30}$$

**$\hat{\mathbf{N}}_{\mathbf{R}} = \mathbf{0}$**

If the equilibrium value of $\hat{N}_R$ is 0, it follows that the equilibrium value of $\hat{Z}_R$ is 0, $f(\cdot) = 1$, $g(\cdot) = 1$ and consequently the equilibrium value of $\hat{N}_T = 1$. The equilibrium values of $\hat{I}$ and $\hat{Z}_T$ come from a solution to

$$0 = i_T \hat{Z}_T\left(1-\hat{I}\right) - \hat{I} \tag{31}$$

$$0 = u + a_T(1-\hat{Z}_T)\hat{I} - (d+e)\hat{Z}_T \tag{32}$$

The second equation indicates

$$\hat{Z}_T = \frac{u + a_T\hat{I}}{a_T\hat{I} + d + e} \tag{33}$$

Back subbing into the first equation

$$0 = \left(i_T \frac{u + a_T\hat{I}}{a_T\hat{I} + d + e}\right)(1-\hat{I}) - \hat{I} \tag{34}$$

The number and nature of the equilibria implied by Eq 34 depends on whether or not there is immigration of viruliferous vectors (*i.e.* $u = 0$ or $u > 0$).

- $\boldsymbol{u = 0}$

  If $u = 0$, which is equivalent to assuming that all the aphids entering the system are non viruliferous, the Eq 34 can be rewritten as

$$0 = \hat{I}\left(\left(\frac{i_T a_T}{a_T\hat{I} + d + e}\right)(1-\hat{I}) - 1\right) \tag{35}$$

  which means there is one root when $\hat{I} = 0$ and one root comes from the solution to

$$\frac{i_T a_T}{a_T\hat{I} + d + e} = \frac{1}{(1-\hat{I})} \tag{36}$$

  After algebraic manipulations, recalling that $R_0^T = \frac{i_T a_T}{d+e}$ when $f(\cdot) = g(\cdot) = 1$, the solution to Eq 36 is

$$\hat{I} = \frac{R_0^T - 1}{R_0^T + a_T/(d+e)} \tag{37}$$

  which, for $R_0^T > 1$, is always in the biologically-meaningful interval [0,1].

- $\boldsymbol{u > 0}$

  If $u > 0$, which is equivalent to assuming that transient aphids can bear the disease from outside the system, $\hat{I} = 0$ is not a solution of Eq 34, so it is acceptable to divide Eq 34 by $\hat{I}$, leading to

$$0 = \left(\frac{i_T u}{\hat{I}(a_T\hat{I} + d + e)} + \frac{i_T a_T}{a_T\hat{I} + d + e}\right)(1-\hat{I}) - 1 \tag{38}$$

$$\frac{1}{1-\hat{I}} = \frac{i_T u}{\hat{I}(a_T\hat{I} + d + e)} + \frac{i_T a_T}{a_T\hat{I} + d + e} \tag{39}$$

$$v(\hat{I}) = b_1(\hat{I}) \tag{40}$$

The function $b_1(\hat{I})$ is: *i)* always positive for $\hat{I} \in (0,1]$; *ii)* always decreasing; *iii)* has $b_1(\hat{I}) \to +\infty$ as $\hat{I} \to 0^+$. The function $v(\hat{I})$ *i)* is always positive for $\hat{I} \in [0,1)$; *ii)* is always increasing; *iii)* has $v(0) = 1$ and has $v(\hat{I}) \to +\infty$ as $\hat{I} \to 1^-$. Taken together the properties of $b_1(\hat{I})$ and $v(\hat{I})$, we can conclude that there is always a single biologically meaningful root with $\hat{I} \in (0,1)$ (irrespective of the values of parameters) (Fig S1A).

**$\hat{\mathbf{N}}_\mathbf{R} = \mathbf{1}$**

If the equilibrium value of $\hat{N}_R$ is 1, it follows that $f(\cdot) = \frac{1}{1+\left(\frac{\nu_1}{\hat{h_R}^{1-\beta_1}\hat{h}^{\beta_1}}\right)^{\alpha_1}} = f$, $g(\cdot) = 1+\left(\frac{\nu_2}{\hat{h_R}^{1-\beta_2}\hat{h}^{\beta_2}}\right)^{\alpha_2} = g$, and the equilibrium value of $\hat{N}_T = \frac{1}{g}$. The equilibrium value of $\hat{I}$, $\hat{Z}_R$ and $\hat{Z}_T$ come from a solution to

$$0 = \Big(i_R\hat{Z}_R + i_T f\hat{Z}_T\Big)(1-\hat{I}) - \hat{I} \quad (41)$$

$$0 = a_R(1-\hat{Z}_R)\hat{I} - m\hat{Z}_R \quad (42)$$

$$0 = u + a_T f\Big(\frac{1}{g} - \hat{Z}_T\Big)\hat{I} - (d+eg)\hat{Z}_T \quad (43)$$

The second and third equations indicate

$$\hat{Z}_R = \frac{a_R\hat{I}}{a_R\hat{I}+m} \quad (44)$$

$$\hat{Z}_T = \frac{ug + a_T f\hat{I}}{g(a_T f\hat{I} + d + eg)} \quad (45)$$

Back subbing into the first equation

$$0 = \left(i_R\frac{a_R\hat{I}}{a_R\hat{I}+m} + i_T f\frac{ug + a_T f\hat{I}}{g(a_T f\hat{I} + d + eg)}\right)(1-\hat{I}) - \hat{I} \quad (46)$$

Again, the number and nature of the equilibria implied by Eq 46 depends on whether or not there is immigration of viruliferous vectors (*i.e.* $u = 0$ or $u > 0$).

- $\boldsymbol{u = 0}$

If $u = 0$, Eq 46 can be rewritten as

$$0 = \hat{I}\left(\left(\frac{i_R a_R}{a_R \hat{I} + m} + \frac{i_T a_T f^2}{g(a_T f \hat{I} + d + eg)}\right)(1 - \hat{I}) - 1\right) \quad (47)$$

which means there is one root when $\hat{I} = 0$ and others come from the solutions to

$$\left(\frac{i_R a_R}{a_R \hat{I} + m} + \frac{i_T a_T f^2}{g(a_T f \hat{I} + d + eg)}\right) = \frac{1}{(1 - \hat{I})}, \quad (48)$$

The exact equilibrium value of $\hat{I}$ can be found by solving Eq 48

$$\left(\frac{i_R a_R / m}{a_R \hat{I}/m + 1} + \frac{i_T a_T f^2/(g(d + eg))}{a_T f \hat{I}/(d + eg) + 1}\right) = \frac{1}{(1 - \hat{I})} \quad (49)$$

$$\frac{R_0^R}{(a_R \hat{I}/m) + 1} + \frac{R_0^T}{a_T f \hat{I}/(d + eg) + 1} = \frac{1}{1 - \hat{I}} \quad (50)$$

After algebraic manipulation Eq 50 can be written as

$$a_2 \hat{I}^2 + a_1 \hat{I} + a_0 = 0 \quad (51)$$

$$p(\hat{I}) = 0 \quad (52)$$

Where

$$a_2 = -\Big(R_0^R \frac{a_T f}{d + eg} + R_0^T \frac{a_R}{m} + \frac{a_R a_T f}{m(d + eg)}\Big) \quad (53)$$

$$a_1 = \Big(\frac{a_T f}{d + eg}(R_0^R - 1) + \frac{a_R}{m}(R_0^T - 1) - (R_0^R + R_0^T)\Big) \quad (54)$$

$$a_0 = R_0^R + R_0^T - 1 \quad (55)$$

Since $a_2 < 0$, when $a_0 > 0$, which corresponds to $R_0^R + R_0^T = R_0 > 1$, the quadratic function $p(\hat{I})$ in Eq 51 has two roots, one positive and one negative. Since $p(1) = a_2 + a_1 + a_0 = -\left(\frac{a_R a_T f}{m(d+eg)} + \frac{a_T f}{d+eg} + \frac{a_R}{m} + 1\right) < 0$ the positive root must have $\hat{I} < 1$ by the intermediate value theorem, thus it is in the biologically meaningful interval $[0, 1]$.

We can conclude that Eq 51 has one biologically meaningful solution, expressed as

$$\hat{I} = \frac{-a_1 - \sqrt{a_1^2 - 4a_2 a_0}}{2a_2} \quad (56)$$

- ***u* > 0**

  If $u > 0$, $\hat{I} = 0$ is not a solution of Eq 46, so it is acceptable to divide Eq 46 by $\hat{I}$, leading to

$$0 = \left( \frac{i_R a_R}{a_R \hat{I} + m} + \frac{i_T f u}{\hat{I}(a_T f \hat{I} + d + eg)} + \frac{i_T a_T f^2}{g(a_T f \hat{I} + d + eg)} \right)(1 - \hat{I}) - 1 \tag{57}$$

$$\frac{1}{1 - \hat{I}} = \left( \frac{i_R a_R}{a_R \hat{I} + m} + \frac{i_T f u}{\hat{I}(a_T f \hat{I} + d + eg)} + \frac{i_T a_T f^2}{g(a_T f \hat{I} + d + eg)} \right) \tag{58}$$

$$v(\hat{I}) = b_2(\hat{I}) \tag{59}$$

  The function, $v(\hat{I})$ has the same properties as before, whereas the function $b_2(\hat{I})$ is: *i*) always positive on the interval $\hat{I} \in (0, 1)$; *ii*) always decreasing; *iii*) has $b_2(\hat{I}) \to +\infty$ as $\hat{I} \to 0^+$. Taken together with the properties of $v(\hat{I})$ this means there is always a single biologically meaningful root with $\hat{I} \in (0, 1)$ (irrespective of the values of parameters) (Fig S1B).

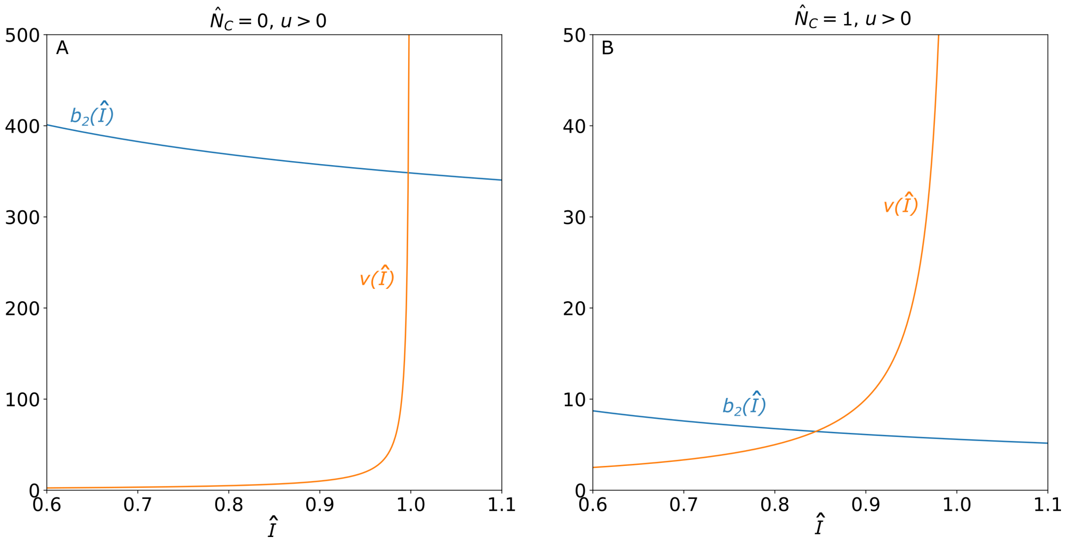


Figure S1: Graphical representation of A) functions $b_1(\hat{I})$ and $v(\hat{I})$ and B) functions $b_2(\hat{I})$ and $v(\hat{I})$. Function $v(\hat{I})$ is the same in A) and B), but values on the y-axis are different to facilitate figure interpretation. The intersection of the two functions identifies the equilibrium value of $\hat{I}$. $\nu = 12.0$, $\pi = 0.1$, other parameters are set to default values.

The model behaviour at equilibrium is summarized in Table A in S2 Text, which enriches Table 2 in the main text. Moreover, for 5000 sets of parameter values randomly selected from the range $[0.8P; 1.2P]$ where $P$ is the default value of each of the parameters, in all cases the result of the numerical simulation matches the results of the mathematical analysis, as well as the stability properties of the equilibria.

Table A: Summary of equilibrium behaviour. The value of the nondimensionalized state variables at the equilibrium are indicated in the forth column. Where, for space limitation, is was not possible to report the equilibrium expression, the reference to the corresponding equation is reported.

| Viruliferous aphids enter the system ($u > 0$) | Resident aphids are present ($q > 0$) | Basic reproduction number (Eqs 24 and 25) | $(\hat{I}, \hat{N}_R, \hat{Z}_R, \hat{N}_T, \hat{Z}_T)$ | Explanation |
|---|---|---|---|---|
| no | no | $R_0 < 1$ | (0, 0, 0, 1, eq. 33) | Transient aphids do not bear the disease from outside the system. Resident aphids are absent, the disease is spread by transient aphids but it does not persist in the system. |
| no | no | $R_0 > 1$ | (eq. 37, 0, 0, 1, eq. 33) | Transient aphids do not bear the disease from outside the system. Resident aphids are absent, the disease is spread by transient aphids. |
| no | yes | $R_0 < 1$ | (0, 1, eq. 44, $\frac{1}{g}$, eq. 45) | Transient aphids do not bear the disease from outside the system. Resident and transient aphids spread the disease, but it does not persist in the system. |
| no | yes | $R_0 > 1$ | (eq. 56, 1, eq. 44, $\frac{1}{g}$, eq. 45) | Transient aphids do not bear the disease from outside the system. Resident and transient aphids spread the disease. |
| yes | no | - * | (eq. 40, 0, 0, 1, eq. 33) | Transient aphids bear the disease from outside the system. Resident aphids are absent, transient aphids spread the disease. |
| yes | no | - * | (eq. 59, 1, eq. 44, $\frac{1}{g}$, eq. 45) | Transient aphids bear the disease from outside the system. Resident and transient aphids spread the disease. |

* The disease is always able to persist, the basic reproduction number is not definable.
