## Supplementary material for "Modelling interference between vectors of non-persistently transmitted plant viruses to identify effective control strategies": S3 Text

### S3 Text. Agricultural practices and disease control

The responses of the incidence of virus infection in plants ($\bar{I}$) and of the basic reproduction number ($R_0$) to variation of plant hosting capacity ($h$), aphid mortality ($\mu$) and infected plant roguing rate ($\rho$) are reported in the figure below. Details on the involved processes are given in the main text.

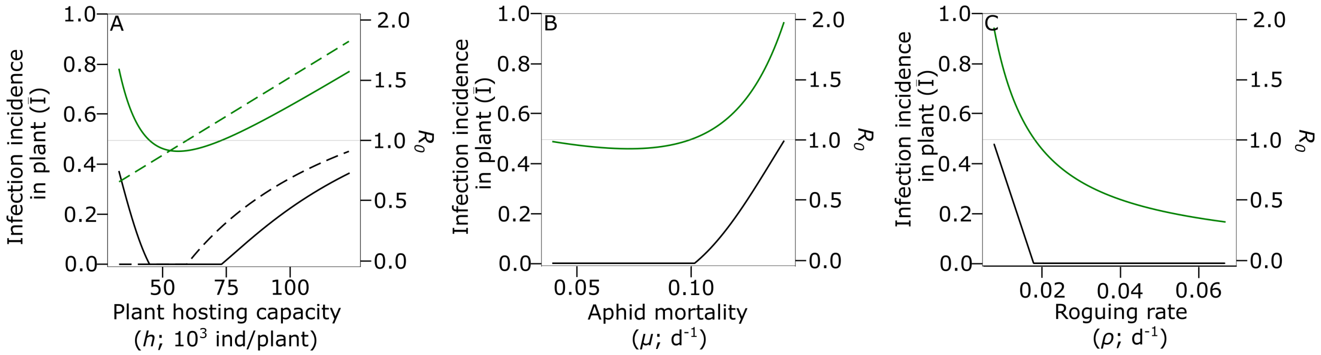


Figure S1: Response of the incidence of virus infection in plant at the equilibrium ($\bar{I}$, in black) and of the reproduction number ($R_0$, in green) to changes in (A) aphids hosting capacity ($h$) under indirect (continuous lines) and direct (dashed lines) interference scenarios; (B) resident aphids mortality ($\mu$); (C) roguing rate ($\rho$).
