## Supplementary material for "Modelling interference between vectors of non-persistently transmitted plant viruses to identify effective control strategies": S4 Text

### S4 Text. Influence of interference parameters on results

We explored the responses of $R_0$ to variation of plant hosting capacity ($h$), aphid mortality ($\mu$) and infected plant roguing rate ($\rho$) under three scenarios of interference strength, corresponding to three values of the interference strength parameter ($\nu = 2$, $\nu = 12$, $\nu = 22$ ). Similarly, we explored the response of $R_0$ to variation of parameters $h$, $\mu$ and $\rho$ under three scenarios of interference curvature, corresponding to three values of the curvature parameter ($\alpha = 0.5$, $\alpha = 1$ and $\alpha = 1.5$). Results are showed in Figs S1 and S2, respectively. The results presented in the main text, corresponding to parameter values $\nu = 12$ and $\alpha = 1$, are shown in the second row of Figs S1 and S2. The responses of $R_0$ to variations of $h$, $\mu$ and $\rho$ are generally qualitative unaffected by the value of $\nu$ and $\alpha$. The only difference is the response of $R_0$ to variation in aphid mortality for $\nu = 2$ (Fig S1B) and $\alpha = 0.5$ (Fig S2B). Here, the response of $R_0$ is monotone, in contrast to the results showed in the main text. In this case, the interference exerted by resident aphids is so low that, even for small value of resident aphids mortality, the virus is mainly transmitted by the transient vector. However, we note that if the values of other parameters were altered, for example to increase the density of resident aphids by increasing the plant hosting capacity $h$, then non-monotonicity would once again be seen. Nevertheless, the general finding that increasing pesticides could be counter productive in reducing NPT viruses is still valid.

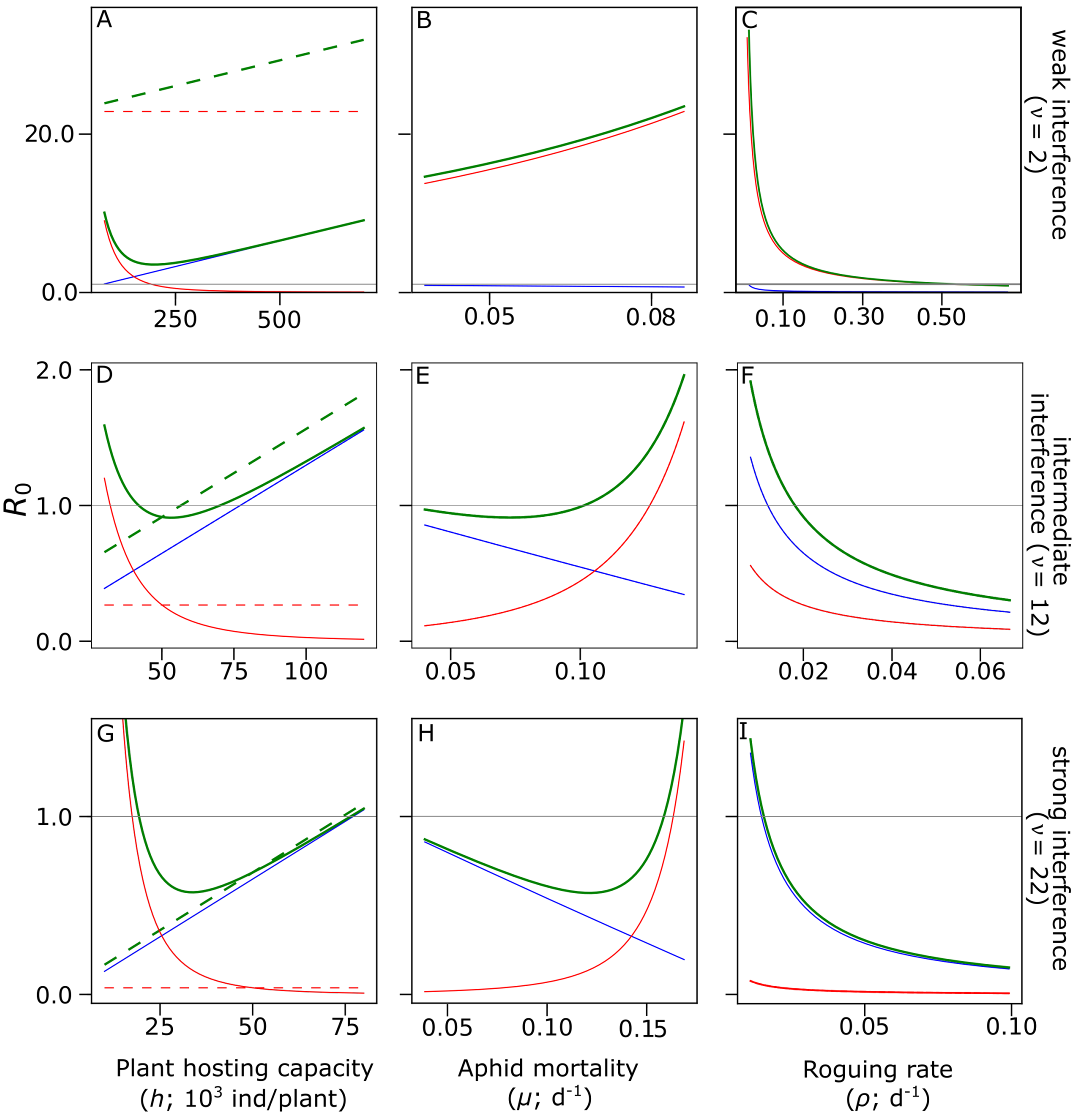


Figure S1: Influence of interference strength parameter ($\nu$) on the response of the basic reproduction number $R_0$ (in bold and green) and its components $R_0^R$ (in blue) and $R_0^T$ (in red) to changes in (A, D, G) plant hosting capacity ($h$) under indirect (continuous line) and direct (dashed line) interference scenarios, (B, E, H) resident aphids mortality ($\mu$), (C, F, I) roguing rate ($\rho$). Note that in (A, D, G) blue continuous and dashed lines overlap, in (C) green and red continuous lines overlap and in (I) green and blue continuous lines overlap.

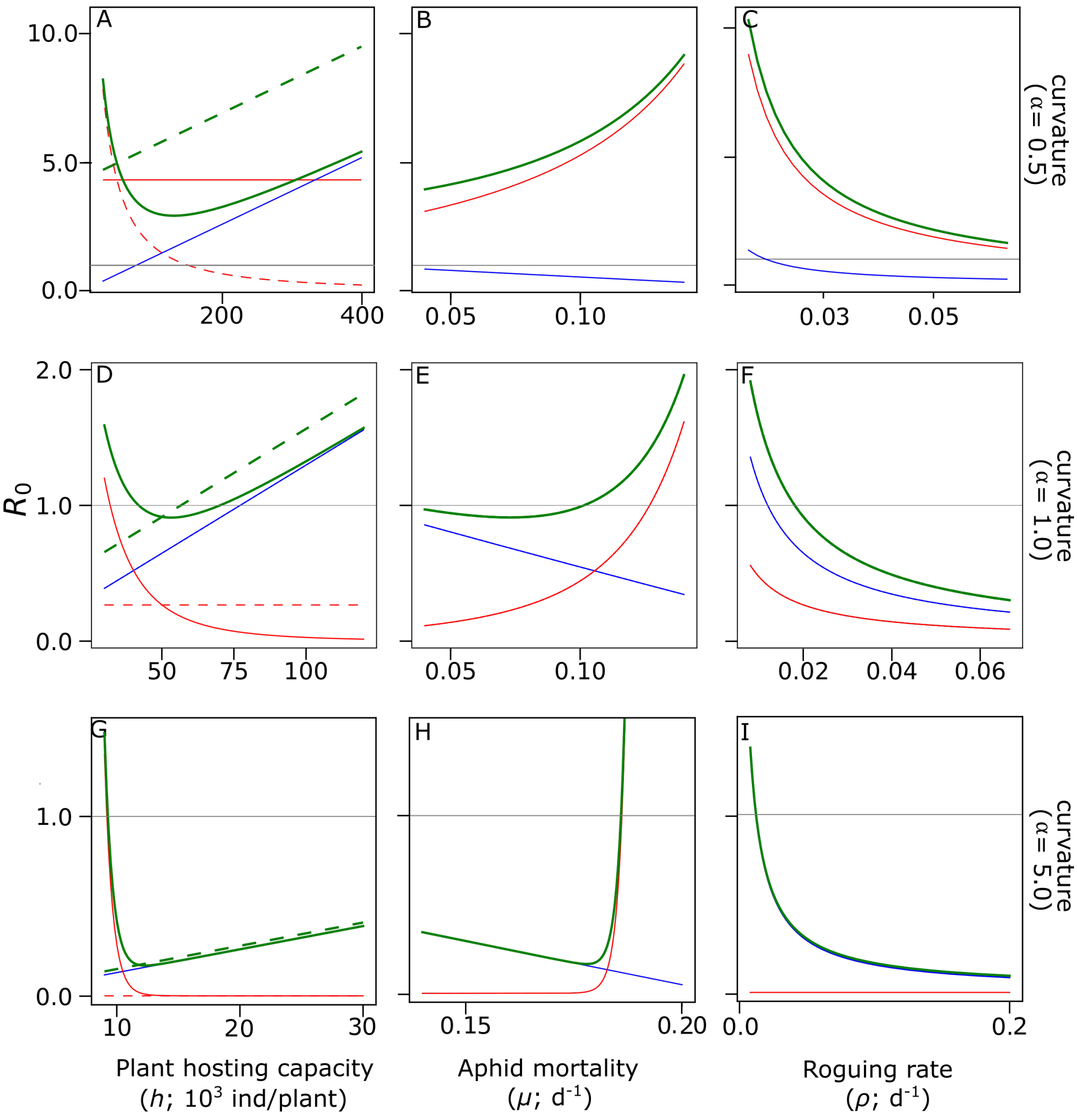


Figure S2: Influence of the interference curvature parameter ($\alpha$) on the response of the basic reproduction number $R_0$ (in bold and green) and its components $R_0^R$ (in blue) and $R_0^T$ (in red) to changes in (A, D, G) plant hosting capacity ($h$) under indirect (continuous line) and direct (dashed line) interference scenarios, (B, E, H) resident aphids mortality ($\mu$), (C, F, I) roguing rate ($\rho$). Note that in (A, D, G) blue continuous and dashed lines overlap, in (G) green continuous and dashed lines and blue continuous and dashed lines overlap for $h > 12000$ ind/plant (green dashed line is slightly moved up to improve visualization), and in (I) green and blue continuous lines overlap.
